# Tracing the Path from 4-Hydroxyphenylpyruvate to the Benzoquinone Ring of Q_6_ and the *p*-aminobenzoate pathway in Yeast

**DOI:** 10.64898/2026.06.24.734323

**Authors:** Maria Jose Valera, Mauricio Mastrogiovanni, Lucia Fernandez del Rio, Eduardo Boido, Juan Carlos Ramos, Eduardo Manta, Eduardo Dellacassa, Rafael Radi, Catherine F. Clarke, Francisco Carrau

## Abstract

Coenzyme Q (ubiquinone, CoQ) is an essential component of the mitochondrial electron transport chain and a major lipid antioxidant in eukaryotic cells. Formation of its benzoquinone ring requires aromatic precursors whose metabolic origin remains incompletely defined. Here, we elucidate the biochemical link between tyrosine metabolism and the synthesis of the benzoquinone head group of coenzyme Q_6_ (Q_6_) in *Saccharomyces cerevisiae* through the 4-hydroxymandelate (4HMA) pathway.

Using isotopic tracing with ^13^C_6_-tyrosine, ^13^C_6_-4-hydroxybenzoate, and ^13^C_6_-*p*-aminobenzoate (*p*ABA), we demonstrate that tyrosine-derived 4-hydroxyphenylpyruvate is converted into 4-hydroxybenzaldehyde via benzoylformate decarboxylation, defining a functional 4HMA pathway in yeast. Chemical inhibition of benzoylformate decarboxylase with methylbenzoylphosphonate led to accumulation of pathway intermediates, which were identified by GCMS. Consistently, mutants lacking *ARO10*, *DLD1*, or *DLD2* exhibited strongly decreased 4-hydroxybenzaldehyde formation.

Despite disruption of the 4HMA pathway, the *p*ABA route from chorismate compensated, demonstrating *S. cerevisiae*’s metabolic flexibility to use *p*ABA or 4-HB and maintain Q_6_ ring biosynthesis. Our results provide a mechanistic framework linking aromatic amino acid metabolism to respiratory quinone biosynthesis in eukaryotes and support the evolutionary conservation of the 4HMA-derived pathway as a source of 4-hydroxybenzoate for Q synthesis in higher organisms.

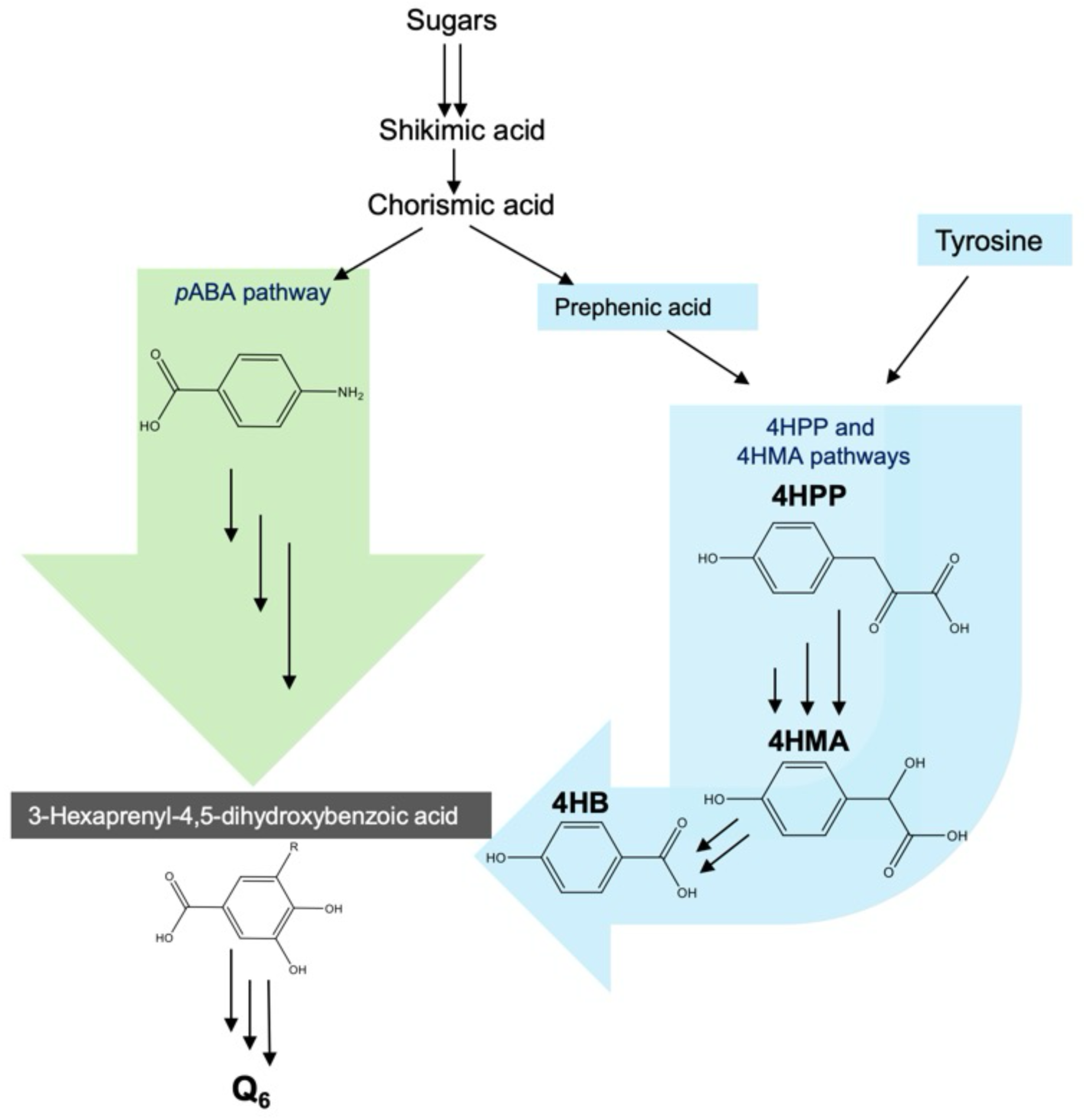
Graphical abstract. 4-hydroxyphenylpyruvic acid (4HPP), 4-hydroxymandelate (4HMA), 4-hydroxybenzoic acid (4HB), *p*-aminobenzoic acid (*p*ABA).

## Introduction

Coenzyme Q (Q) was discovered to be a redox-active, lipophilic compound that is essential for oxidative phosphorylation in eukaryotes (1), and in a wide array of species within the diverse bacterial phyla Pseudomonadota (2). The universal aromatic ring precursor leading to all Qn isoforms is 4-hydroxybenzoic acid (4HB), present in organisms ranging from *E. coli* and plants to humans (3, 4). The aromatic ring precursor is linked to a long hydrophobic polyisoprene tail, with a varying number of isoprene units depending on the species (4, 5). For example, *Schizosaccharomyces pombe* synthesize Q_10_, *Saccharomyces cerevisiae* synthesize Q_6_, *E. coli* synthesize Q_8_, *C. elegans* and rats synthesize Q_9_, and most mammals, including humans, synthesize Q_10_ (6, 7).

Qn biosynthesis represents a metabolic convergence between aromatic amino acid metabolism and the mevalonate pathway. While the isoprenoid tail is derived from the mevalonate pathway, 4HB is derived from tyrosine catabolism and/or the chorismate pathway in yeasts and bacteria (8). *p*ABA derived from chorismate can also serve as a Q ring precursor in the yeasts *S. cerevisiae* (9, 10), *Schizosaccharomyces pombe* (11), and in *Hanseniaspora vineae* (Valera et al. submitted). Payet et al (8) demonstrated that the aminotransferases encoded by the *ARO8* and *ARO9* genes interconvert tyrosine and 4-hydroxyphenylpyruvate (4HPP) (see Fig. 1), and catalyze the first step of the phenylpyruvate pathway (12, 13). Payet et al. (8) and Stefely et al., (14) identified that the dehydrogenase encoded by *HFD1* was responsible for the final step, the conversion of 4-hydroxybenzaldehyde (4HBz) to 4HB. The *hfd1* yeast mutant can be rescued by exogenous supplementation with 4HB, underscoring the essential role of this step in the supply of 4HB. Although some studies have suggested pathways to 4HB synthesis in yeast (8, 13–17), the low levels of intermediates and enzyme activities, redundancies, and the existence of *p*ABA as an alternative precursor in *S. cerevisiae* have hindered attempts to demonstrate this definitively (12, 16).

**Fig 1.**
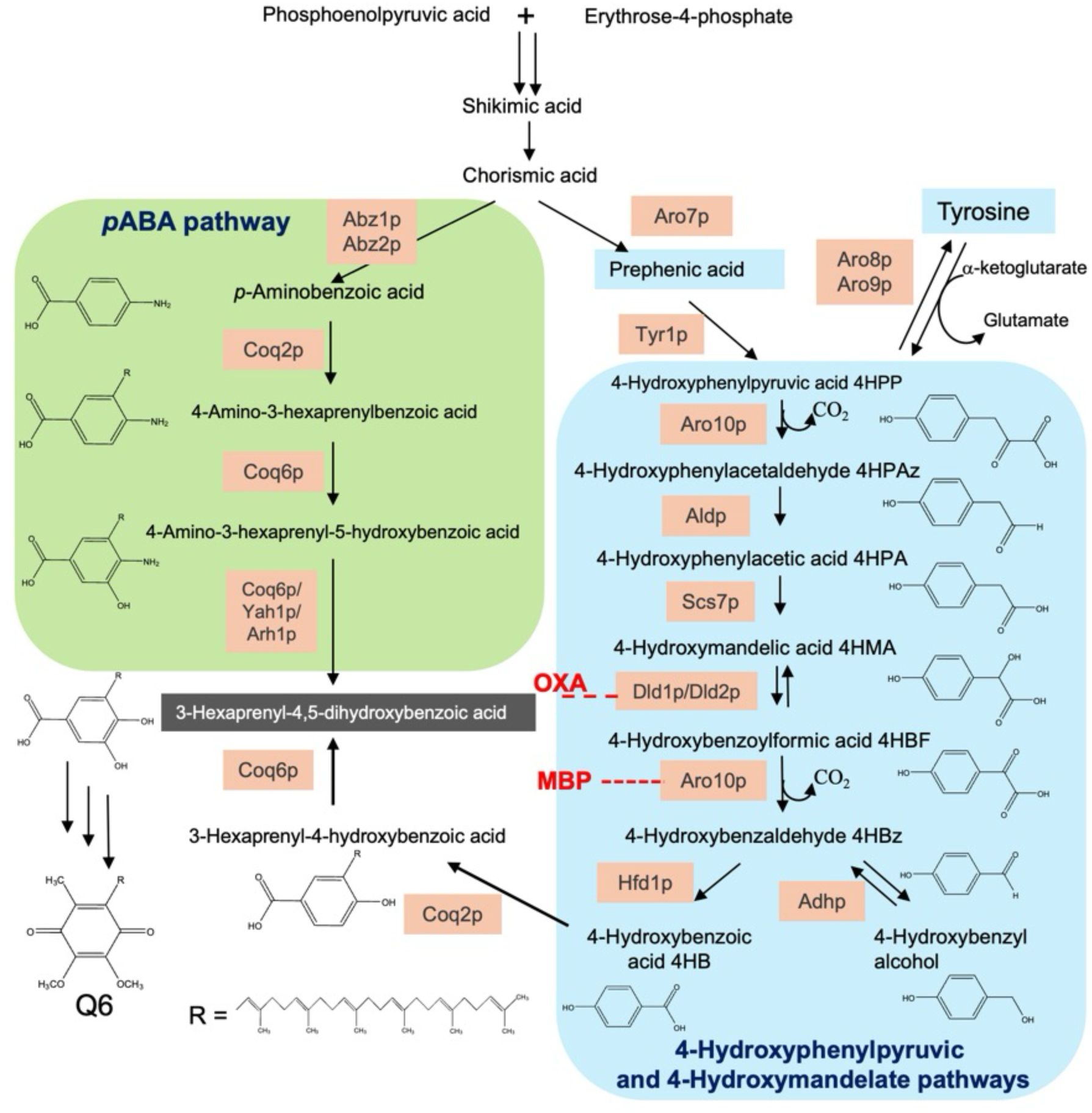
Two alternative biosynthetic pathways for Q6 formation in *S. cerevisiae* utilize either *p*ABA or tyrosine as aromatic ring precursors. The link between 4-hydroxyphenylpyruvic acid and 4-hydroxybenzaldehyde is completed via the 4-hydroxymandelate (4HMA) pathway. Identification of the key intermediates by GC-MS and HPLC-MS was facilitated by inhibition of 4-hydroxybenzoylformic acid decarboxylase (the gene product of *ARO10*) using the methylbenzoylphosphonate (MBP) analogue, and by oxamate (OXA) at the 4-hydroxymandelic acid dehydrogenase step (encoded by *DLD1/DLD2*) (both shown in red). As described in the text, *aro10* mutant trials and *p*ABA compensation of Q6 synthesis confirmed this pathway. 4-hydroxyphenylpyruvic acid (4HPP), 4-hydroxyphenylacetaldehyde (4HPAz), 4-hydroxyphenylacetic acid (4HPA), 4-hydroxy mandelate (4HMA), 4-hydroxybenzoylformic acid (4HBF), 4-hydroxybenzaldehyde (4HBz), 4-hydroxybenzoic acid (4HB), *p*-aminobenzoic acid (*p*ABA).

Our studies of the 4HB biosynthesis pathways in *H. vineae*, showed that it produced benzenoid intermediates at concentrations up to two orders of magnitude higher than *S. cerevisiae*. This characteristic made it easier to delineate these reactions originating from tyrosine. Additionally, we found that limiting availability of assimilable nitrogen in the growth medium significantly increased benzenoids production by *H. vineae* (15). We proposed that *H. vineae* utilize 4-hydroxymandelate (4HMA) as an intermediate in the conversion of 4HPP to 4HBz, and that several other intermediate compounds were also involved in the biosynthesis of 4HBz (12). The use of a synthetic medium with very low assimilable nitrogen allowed us to increase the benzenoid concentration to several hundred of micrograms per liter in *H. vineae*. These experiments enabled us to investigate the synthesis 4HBz in a synthetic medium containing ^13^C-tyrosine. When we tested this synthetic medium with *S. cerevisiae*, we also increased benzenoids to levels above the detection limit of the GC-MS method (12). More recently, we showed that *ARO10* encoded an enzyme with benzoylformate decarboxylase function in *S. cerevisiae* (18). These results were consistent with *H. vineae* transcriptome analysis, which showed expression of the eight genes proposed for the complete putative pathway in the first four days of fermentation (12). However, determination of the Q_6_ content was necessary to verify the importance of this putative pathway.

Interestingly, several independent studies support our proposed pathway (12). In yeast mutants, Q_6_ biosynthesis can be restored by supplementation with 4HMA (16), and the formation of 4HMA has been demonstrated in human cells through ^18^O₂ incorporation experiments (19). Although Robinson et al. (16) did not identify the pathway intermediates, they showed that both 4-hydroxyphenylacetaldehyde (4HPAA) and 4HMA rescued mutant strains with respiratory growth defect (*Δaro2 Δaro8 Δaro9)* in *Saccharomyces*. Consistently, Banh et al. (19) detected ^18^O₂-labeled 4HMA in human cell lines derived from multiple tissue types. Furthermore, Shi et al. (20) demonstrated that oral administration of 4HMA in mice results in its metabolic incorporation in vivo and restores Q_10_ levels in animals with Q_10_ deficiency and mitochondrial encephalopathy. Together, these findings suggest that the 4HMA pathway may contribute to Q biosynthesis in both yeast and mammalian cells. However, the sequence of intermediates in this pathway remains unresolved and will require confirmation through ^13^C metabolic tracing by GC-MS or HPLC-MS and detailed Q_6_ analysis (13, 17).

In the current study, we used a synthetic medium containing low levels of ammonia and amino acids, along with the labelled precursors: ^13^C_6_-Phe, ^13^C_6_-Tyr and ^13^C_6_-*p*ABA. This allowed us to detect the intracellular and extracellular benzenoids in agreement with the measured compounds in *H. vineae* trials (Valera et al. submitted), and to detect and quantify the Q_6_ produced by *S. cerevisiae* under log phase growth and fermentative conditions. Interestingly, our results suggest that the *p*ABA pathway in *Saccharomyces* might be the main source of the Q_6_ benzoquinone head.

## Results

Several genes have been proposed to be involved in 4HBz biosynthesis, the precursor of 4HB (Table 1 and Fig. 1). Different experiments were performed to analyze the implication of these gene products in the biosynthesis of 4HBz and Q_6_. Two main conditions were defined for these experiments: fermentation and growth. Fermentation experiments were characterized by long duration (5 to 12 days) in a medium with high sugar concentration (200 g/L) to favor anaerobic metabolism and low nitrogen concentration (100 mgN/L of yeast assimilable nitrogen). In log-phase growth experiments, sugar concentrations were low (20 g/L of glucose) the cultures were well aerated for shorter times (between 16 and 48 hours) promoting aerobic metabolism and nitrogen concentration was also 100 mg N/L of yeast assimilable nitrogen.

**Table 1.** ***S. cerevisiae* genes involved in the 4-HBz synthesis and their role in the pathway**

| Gene | Protein | Subcellular localization | Main enzymatic activity* | Link to the 4-hydroxymandelic acid pathway* | Validation |
| --- | --- | --- | --- | --- | --- |
| ***ARO10*** | Aro10p | Cytoplasm | Aromatic α-keto acid decarboxylase. Benzoylformate Decarboxylase function. | Generates aromatic aldehydes from 4-hydroxyphenylpyruvate; defines metabolic fluxes that may compete with or divert precursors from 4-hydroxymandelic acid formation. Inhibited by MBP. | Deleted mutant of Sc inhibits 95% 4-HBz formation. It catalyzes two decarboxylation steps shown in Fig. 1. |
| ***DLD1*** | Dld1p | Mitochondria | D-lactate dehydrogenase (FAD-dependent) | Potential oxidation of aromatic α-hydroxy acids; may participate in mitochondrial metabolism of 4-hydroxymandelic acid or related intermediates. Inhibited by Oxamate. | Deleted mutant of Sc inhibits 90% 4-HBz formation. |
| ***DLD2*** | Dld2p | Cytoplasm | D-lactate dehydrogenase (NAD<sup>+</sup>-dependent) | Candidate enzyme for cytosolic redox interconversion of 4-hydroxymandelic acid; contributes to α-hydroxy acid redox balance. Inhibited by Oxamate. | Deleted mutant of Sc inhibits 50% 4-HBz formation. |
| ***ALD2*** | Ald2p | Cytoplasm | Aldehyde dehydrogenase (NAD<sup>+</sup>-dependent) | Oxidation of aromatic aldehydes derived from 4-hydroxymandelic acid (e.g., 4-hydroxybenzaldehyde) to phenolic acids. |  |
| ***ALD6*** | Ald6p | Cytoplasm | Aldehyde dehydrogenase (NADP<sup>+</sup>-dependent) | Conversion of phenolic aldehydes to aromatic acids with NADPH generation; contributes to the formation of 4-hydroxybenzoate. |  |
| ***SCS7*** | Scs7p | Endoplasmic reticulum | Very-long-chain fatty acid hydroxylase | Indirect role: modulates membrane integrity and oxidative status, potentially influencing mitochondrial and aromatic metabolic processes. |  |

### Deletion mutants of *S. cerevisiae* mimic inhibition by MBP and OXA in 4-HBz biosynthesis

Select diploid homozygote mutants from the *S. cerevisiae* knockout collection were tested due to their predicted involvement in the formation of extracellular 4HBz after 12 days of fermentation: *dld1Δ, dld2Δ,* and *aro10Δ*. Significant decreases of 4HBz were observed in each of the three deletion mutants (Fig 2A).

**Fig 2.**
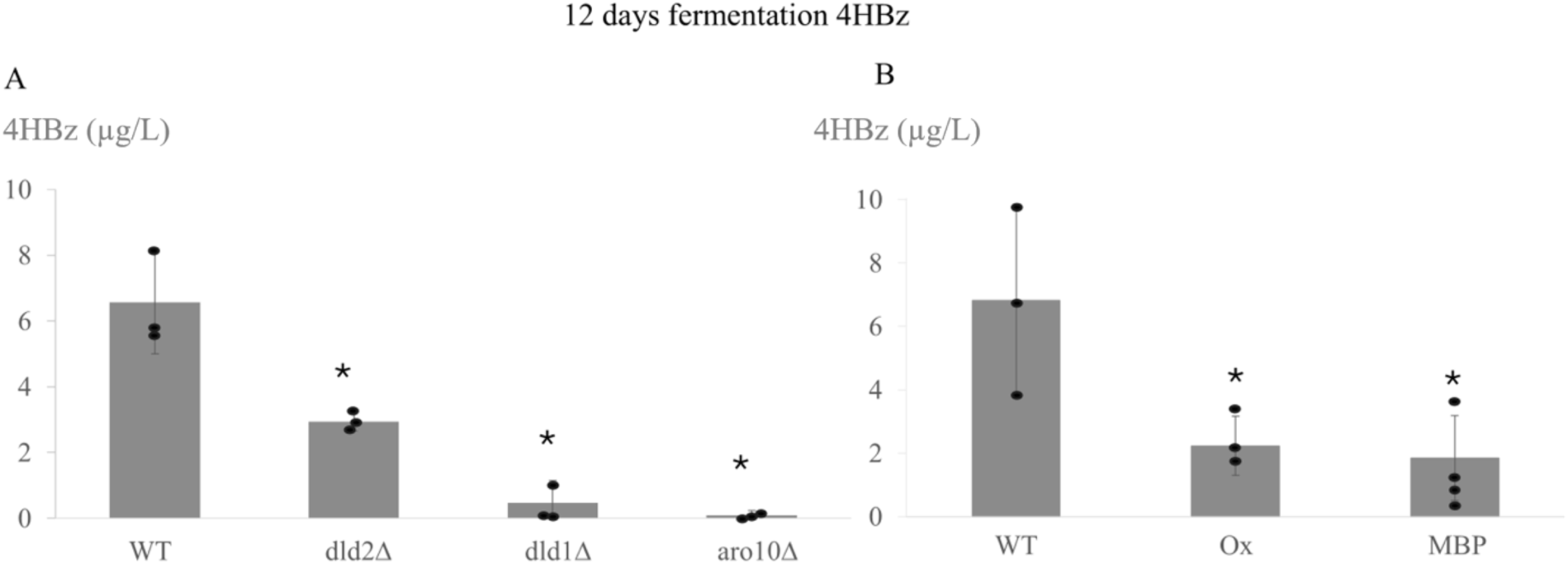
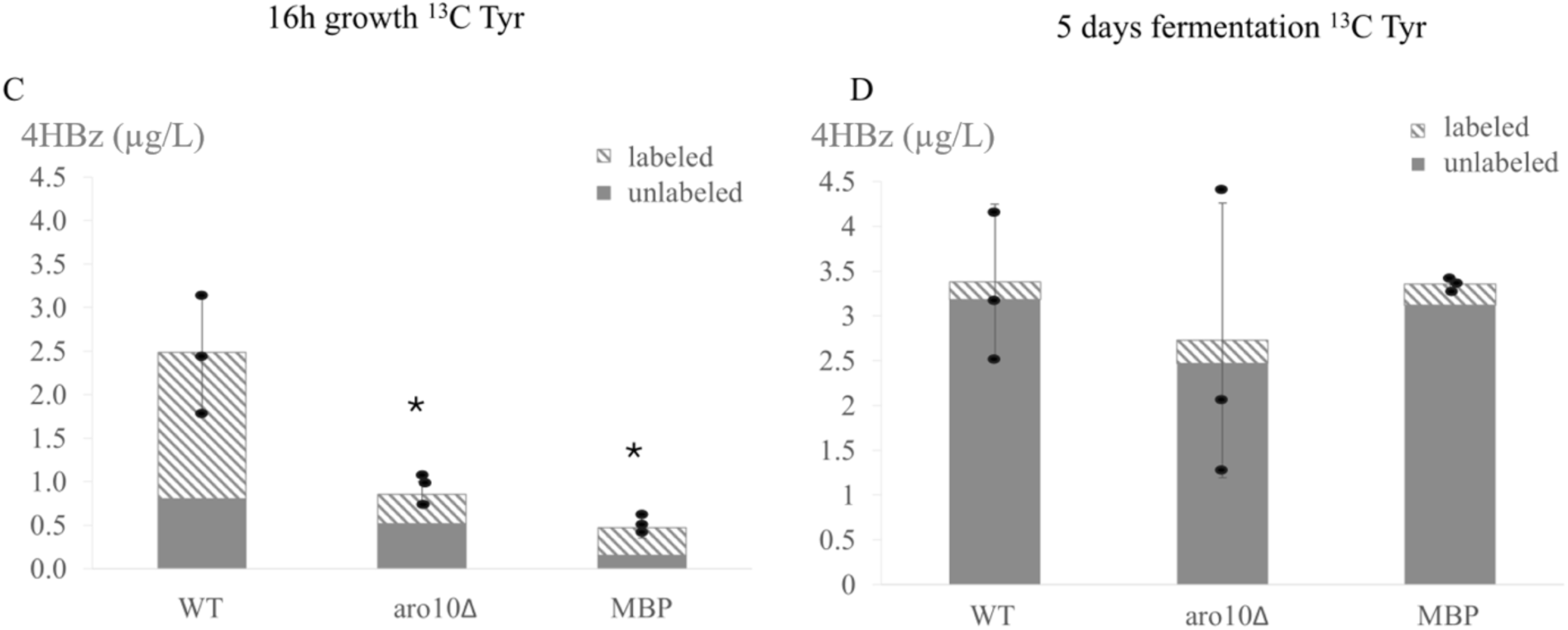
Treatment of *S. cerevisiae* with MBP or OXA inhibitors, or gene deletions of *ARO10*, *DLD1*, or *DLD2* decrease the formation of 4-hydroxybenzaldehyde. In panels A and B, 4HBz was quantified after a period of 12 days of fermentation. In panel C, cultures of WT, *aro10Δ*, or MBP treatment of WT were incubated for 16 h of growth in the presence of ^13^C6-Tyr. In panel D, cultures were conducted as for panel C, except under fermentation conditions for five days. Growth and fermentation experiments were performed using independent triplicate samples and error bars express standard deviation.

Morever, methylbenzoylphosphonate (MBP), an analogue of benzoylformate (21), was used to inhibit benzoylformate decarboxylase (Aro10p), the key enzyme in the 4-hydroxymandelate pathway. In a low assimilable nitrogen synthetic medium (100 mg N/L) (12), 4HBz accumulated in *S. cerevisiae* as detected by GC-MS (Fig. 2B). MBP inhibition of 4HBz formation has been demonstrated previously in *H. vineae*, which produces benzenoids more efficiently than *S. cerevisiae* (12). Furthermore, oxamate (OXA) is known to inhibit D-lactate dehydrogenases (Dld1p and Dld2p) of yeast (22). Treatment with OXA clearly and significantly inhibited the formation of 4HBz in the same way that MBP did (Fig. 2B).

Using MBP and OXA, which inhibit known chemical mechanisms, together with the three deletion mutants of *S. cerevisiae,* we confirmed the involvement of these three genes in the biosynthesis of 4HBz suggesting the following steps in the pathway of the synthesis of 4HB after the formation of 4-hydroxyphenylpyruvate from tyrosine, were the formation of 4-hydroxyphenylacetaldehyde, 4-hydroxyphenylacetate and 4-hydroxymandelate, in a manner similar to that observed in *H. vineae* yeast (Fig. 1) (Valera et al. submitted).

In log-phase growth conditions (16 hours after inoculation), a high proportion of 4HBz appears labeled from ^13^C_6_-Tyr incorporated during exponential growth (Fig. 2C), confirming the strong inhibition of this synthesis in the *aro10Δ* mutant and the MBP treatment. However, under fermentative conditions at day 5, no differences in concentration are detected between the *S. cerevisiae* WT and either the *aro10Δ* mutant or MBP (Fig. 2D). In contrast, at day 12 of fermentation (Fig. 2B), a significantly higher concentration of extracellular 4HBz was accumulated in the WT strain compared with the mutant and MBP treatments, indicating that longer fermentation time substantially affects the process. The WT strain practically duplicates 4HBz accumulation between 5 days (Fig. 2D) and 12 days of fermentation (Fig. 2B), while the mutant and the MBP treatment were stuck during this period.

This effect might be explained by considering extracellular 4HBz as a waste product of secondary metabolism that has flavor impact but is not necessarily utilized by the cell (23). Also, the proportion of 4HBz synthesized from ^13^C_6_-Tyr decreases under fermentation conditions compared with growth, despite similar total amounts. This may be due to the extended duration of fermentation during which ^13^C_6_-Tyr becomes depleted.

### Effect of MBP and *aro10* deletion on Q_6_ synthesis and the alternative *p*ABA pathwa

Although, we confirmed significant effects with *aro10Δ* mutant and MBP under fermentation and log phase growth conditions with labelled ^13^C_6_-Tyr over 4HBz biosynthesis, no significant effects were found in Q_6_ formation, even in starved *p*ABA conditions without folate (Fig 3A and 3B). However, a trend toward lower values was detected in the mutant strain and in the presence of the inhibitor. Incorporation percentages of ^13^C_6_-Tyr into Q_6_ are increased during log phase growth conditions (Fig 3C) as compared to fermentation culture conditions (Fig 3D). The fact that the content of Q_6_ is not significantly decreased in the *aro10Δ* mutant or in the MBP treatment compared with WT strain leads us to question whether Q_6_ formation is being compensated by the alternative *p*ABA pathway discovered in *S. cerevisiae* (9, 10).

**Fig 3.**
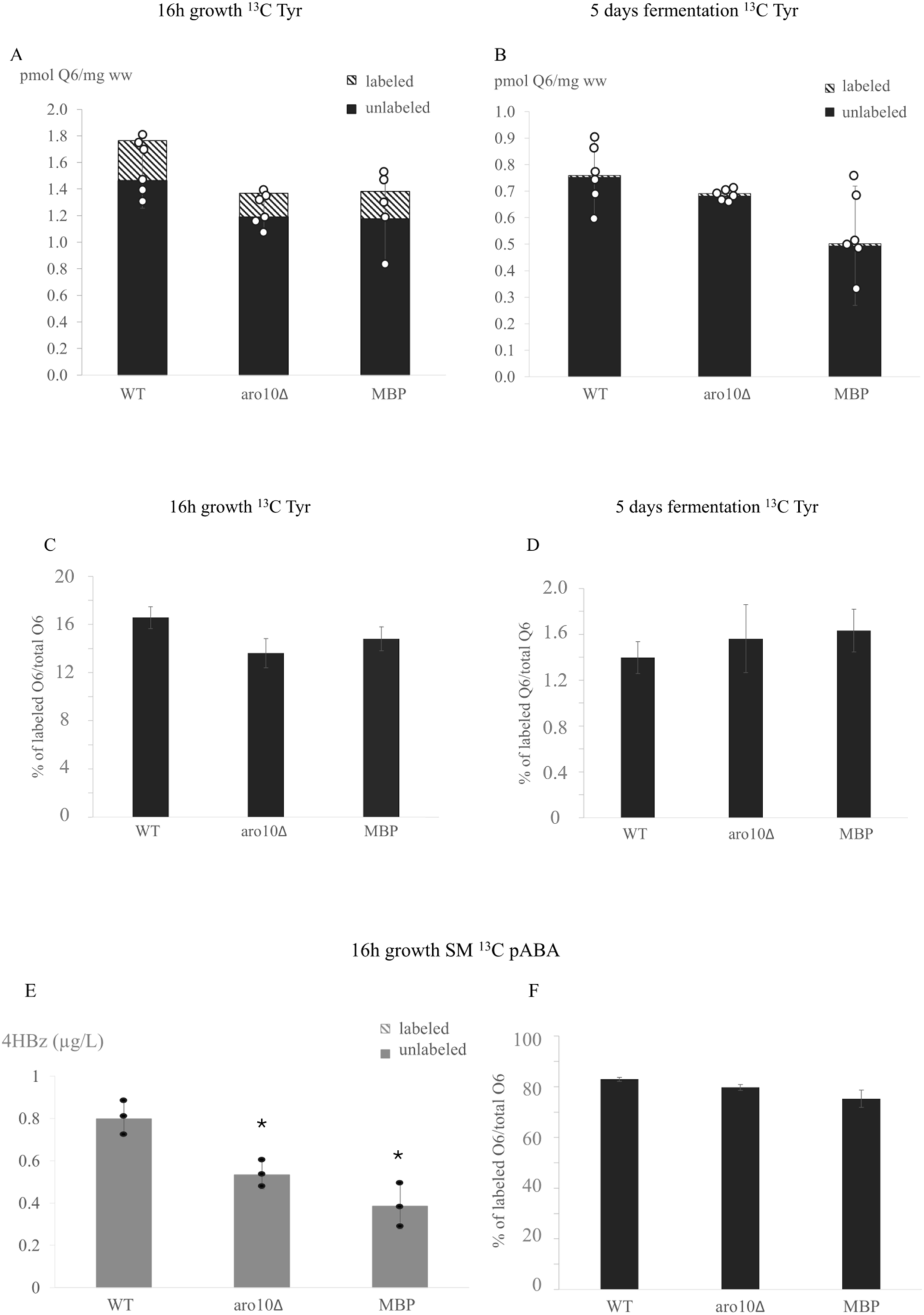
Synthesis of Q6 in fermentation (5 days) and growth (16h) conditions using *S. cerevisiae* WT, *aro10Δ* and MBP. Panel A and B show the quantification of Q6 adding ^13^C6-Tyr. Panel C and D compare percentages of labeled Q6 determined in the treatments. Panel E presents the results of synthesis of 4HBz with ^13^C6-*p*ABA added during exponential growth and F the percentage of labeled of Q6 in these conditions. Growth and fermentation experiments were performed using independent triplicate samples and error bars express standard deviation.

Therefore, we conducted a similar experiment under growth conditions to determine the potential origin of Q_6_ from labeled *p*ABA. The results shown in Fig. 3E and 3F clearly demonstrate that the amount of labeled ^13^C-Q_6_ increased in the presence of *p*ABA compared to the ^13^C_6_-Tyr experiments. Interestingly, the labeled percentages of WT *S. cerevisiae* show that *p*ABA is used as the precursor of the benzenoid core of Q_6_ (Fig. 3F), with a usage rate at 16 hours of growth of about five times higher than that from tyrosine (Fig. 3C). The ability to use *p*ABA for synthesis of the benzoquinone ring of Q_6_ may compensate for the inhibited 4HMA pathway of these treatments. As expected, 4HBz is not labeled when ^13^C_6_-*p*ABA is added in exponential phase, but the inhibitor MBP still decreased significantly the amount of this product as well as the mutant lacking Aro10p (Fig. 3E).

### Synthesis of Q_6_ under growth or fermentation conditions

As there were differences between our experiments involving fermentation and growth, we measured Q_6_ formation under both conditions to more accurately determine the quantitative differences resulting from yeast metabolism. Experiments related to Q synthesis in yeasts are usually conducted during the exponential growth phase, and we did not find studies comparing synthesis under fermentation conditions. Briefly growth conditions are in the first 24 hours of inoculation of the synthetic medium described in methods where a more aerobic situation and lower sugars (20g/L) are present and the exponential phase of growth occurred. Fermentation conditions are determined where the stationary phase occurred, samples are taken between 5 and 12 days, where indicated as the culture medium is rich in sugars (200 g/L) with a very reductive environment due to the CO_2_ production. Considering the exponential growth phase between 16 and 24 hours and the highly active fermentation process on day 5 and near the end of fermentation on day 12 (with still about 10% of sugars present in the medium), we selected these time points to measure Q_6_. The results of these experiments are shown in Fig. 4.

**Fig. 4.**
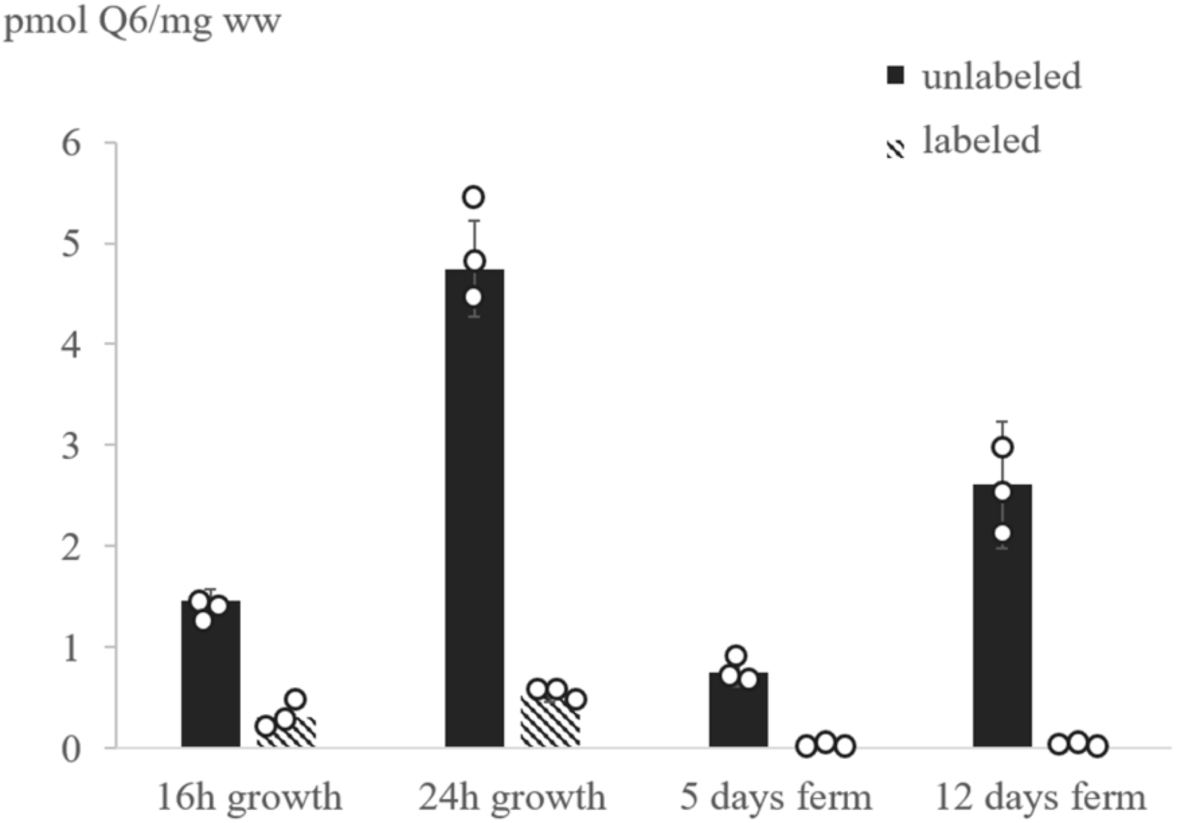
Comparing fermentation with log phase growth, there are significant differences in Q6 synthesis. In growth conditions an aerobic situation is present, with a significant increase in Q6 synthesis between 16 and 24 hours. The latter marks the end of exponential growth in these conditions. In fermentation conditions, the highest concentration of Q6 can be observed on day 12 but lower concentration of Q is detected compare with growth conditions. Labelled Q6 was with ^13^C_6_-Tyr.

Interestingly, when comparing the best studied conditions for Q_6_ synthesis in growth and fermentation, yeast cells in growth conditions synthesized Q_6_ in 24 hours, producing approximately five times more than cells in fermentation conditions over five days. A higher percentage of Q_6_ is labelled under growth conditions, probably due to the shorter growth time compared to fermentation, where ^13^C_6_-Tyr was consumed and supplied internally by synthesis from sugars.

## Discussion

We show that *p-*hydroxybenzenoids biosynthesis in *S. cerevisiae* proceeds via a tyrosine-derived 4-hydroxymandelate (4HMA) pathway that converges on 4-hydroxybenzaldehyde (4HBz), which is then converted to 4-hydroxybenzoate (4HB) and ultimately into coenzyme Q_6_. Genetic and biochemical perturbations—*aro10Δ* mutant and inhibition of Aro10p by methylbenzoylphosphonate (MBP), *dld1Δ/dld2Δ* mutants and inhibition of D-lactate dehydrogenases by oxamate (OXA)—substantially reduced extracellular 4HBz accumulation, supporting roles for Aro10p, Dld1p and Dld2p in steps downstream of 4-hydroxyphenylpyruvate (Fig. 1). Aro10p acts in vivo as a benzoylformate decarboxylase confirming our previous proposal for yeast (18). Deletion of *aro10Δ* and treatment with MBP strongly impaired ^13^C_6_-Tyr labeling of 4HBz during aerobic growth, consistent with a major flux through the 4HMA route under these conditions. The transamination of tyrosine to 4-hydroxyphenylpyruvate and its subsequent conversions via the 4-hydroxymandelate pathway to 4HBz are supported by our mutant and inhibitor data (Figs. 1–2). *S. cerevisiae* efficiently converts 4HBz to 4HB via Hfd1p, providing the headgroup precursor for Q_6_ as was demonstrated previously (8). The proposed intermediate reactions—decarboxylation of 4HPP, oxidation of 4HPA, α-hydroxylation of 4HPAA, oxidation of 4HMA, and decarboxylation of 4HBFA to yield 4HBz—are chemically consistent and are corroborated by three mutant phenotypes, MBP and OXA enzyme inhibitors, metabolite quantification, and *p*ABA compensation of Q_6_ synthesis. These results agree with our report of the yeast *H. vineae* studies, that shows significantly higher intermediates concentrations than *S. cerevisiae* (Valera et al. submitted). Although in this work we couldn’t quantify 4HPAA in *H. vineae*, the fact that addition of it strongly rescued *Δaro2/Δaro8/Δaro9* mutant of *Saccharomyces* as was reported (16), agree with our 4HMA pathway steps (Fig. 1).

We observed condition-dependent differences in Q_6_ synthesis: aerobic (log-phase) cultures showed higher Q*_6_* production compared with long-term fermentation cultures. To our knowledge, previous studies about this phenomenon have not been reported, and the results obtained indicate that oxygen availability, redox state, and growth phase strongly influence Q_6_ concentrations.

The ability of *p*ABA to label Q_6_ more efficiently than tyrosine under growth conditions suggests that the 4-hydroxymandelate pathway needs to incorporate ^13^C_6_-tyr and must metabolize it to 4HB before its use. This concept agrees to the similar incorporation efficiency to *p*ABA of the ^13^C_6_-4HB to Q_6_ found in the yeast *H. vineae* (Valera et al. submitted).

Elucidating the 4HMA pathway in yeast creates new opportunities to explore this pathway in animals. Furthermore, our results encourage the exploration of this pathway in humans, given that 4HMA has been recently detected in human cells and oral supplementation with 4HMA increases Q synthesis. According to a few reports that had detected 4HMA in human cells, supplementation of 4HMA in mice and humans showed recovery of Q_9_ and Q_10_, (20). Some homologue genes found in yeast and humans, such as *HFD1* for the synthesis of 4HB (8), also suggest the same pathway is used by mammalian cells. These experiments confirm that Aro10p of *S. cerevisiae* is involved in 4HBz biosynthesis as a benzoylformic acid decarboxylase, as previously stated by us for *H. vineae* (18).

In summary, the 4-hydroxymandelate pathway in yeast reveals an endogenous tyrosine-derived route to 4HB that contributes to Q_6_ biosynthesis and highlights metabolic plasticity through the alternative *p*ABA pathway, to our knowledge only reported in yeast until now. Given reports of 4HMA detection in mammalian cells and conservation of homologous enzymes (e.g., HFD1p), these findings motivate investigation of analogous pathways in animals and their potential nutritional or therapeutic implications.

## Materials and Methods

### Yeast strains and cultures

Homozygotic diploid double mutant strains *S. cerevisiae* Δ*aro10*, Δ*dld1,* and Δ*dld2* from the Yeast Knockout Collection (24) were supplied by Dharmacon (Lafayette, CO). *S. cerevisiae* BY4743 was used as the control strain for mutant and inhibition experiments. *S. cerevisiae* BY4743 strains were grown in YPD medium (2% glucose, 2% peptone, 1% yeast extract). For growth of the *S. cerevisiae* mutant strains, YPD medium was supplemented with 200 µg/mL of Gentamycin G418 (Acros-Fischer, Belgium).

### Fermentation and growth conditions

The fermentation medium was prepared as previously described (25), with the amino acid content modified according to experimental design. All fermentations were performed with 100 mg N/L of yeast assimilable nitrogen (YAN level, i.e. the sum of the amino acid and ammonium concentrations not including proline) supplemented with 125 mg/L of histidine, 500 mg/L of lysine, 150 mg/L of uracil and 500 mg/L of leucine. The content of each amino acid, expressed in mg/L, is detailed in Table S1. The pH of each medium was adjusted with HCl to a final pH of 3.5. Equimolar concentrations of glucose and fructose were added to reach 200 g/L and mixed vitamins and salts were added as previously described (25) without pABA and folate. Ergosterol (10 mg/L) was the only lipid supplement added. In the fermentation medium all the tyrosine was labelled phenyl ^13^C6-L-tyrosine (Merck Group, Germany)

Inocula were prepared in the same fermentation medium, containing 100 mg N/L of YAN and the same specific amino acid concentrations previously detailed. Erlenmeyer flasks were incubated for 12 h in a rotary shaker at 150 rpm and 25 °C. Fermentations with *S. cerevisiae*, due to their reduced production of benzenoids, were conducted in 500 mL Erlenmeyer flasks with 250 mL of medium to facilitate detection. All the fermentations were closed with cotton plugs to simulate microaerobic conditions (26). The inoculum size was 1x10^5^ cells/mL in the final medium for all strains. Static batch fermentation conditions were conducted at 20 °C and each experiment was performed in triplicate. Fermentation activity was measured as CO2 weight loss and expressed in g/100 mL. Samples were taken for GC-MS analyses following either six or 12 days of the fermentation process.

The respiration medium was prepared with the amino acid content detailed in Table 2. Glucose concentration was 20 g/L and no fructose was added. The pH of the medium was adjusted with HCl to a final pH of 3.5. Equimolar concentrations of glucose and fructose were added to reach 200 g/L and mixed vitamins and salts were added as previously described (25) without pABA and folate. Ergosterol (10 mg/L) was the only lipid supplement added. *S. cerevisiae* strains were conducted in 125 mL Erlenmeyer flasks with 30 mL of medium. The inoculum size was 1x10^6^ cells/mL in the final medium. Shaken cultures were maintained at 25 °C and each experiment was performed in triplicate. In the respiration experiments either labelled tyrosine phenyl ^13^C6-L-tyrosine or labelled *p*ABA phenyl ^13^C6-p-Aminobenzoic acid (Merck Group, Germany) were added in mid-exponential phase (OD620 ≈ 0.5) to a final concentration of 5µg/mL. The entire volume of each flask was subjected to centrifugation at 8,000 rpm for 10 min to separate the cells from the extracellular medium, after which the extracellular medium was filtered using 0.45 μm pore membranes.

### Inhibition experiments by MBP and oxamate

MBP was synthesised according to the method of Brandt et al.(27). Briefly, 4.75 mL of trimethylphosphite (5 g, 40.3 mmol) was added in a dropwise fashion to a stirred volume of benzoyl chloride (5.66 g, 40.3 mmol) under nitrogen at 0 °C. After the addition was completed, the reaction mixture was stirred for 30 min at 0 °C and then allowed to warm to room temperature until no benzoyl chloride was observable using thin layer chromatography. The resulting yellowish oil was purified in a chromatography column yielding 7.1 g (82%), ^1^H NMR (400 MHz, CDCl3, 0.5% TMS): δ = 8.24 (d, J 7.5 Hz, 2H), 7.53 (t, J 7.5 Hz, 1H), 7.494 (t, J 7.5 Hz, 2H), 3.910 (d, J 10.5 Hz, 6H). Dimethyl benzoylphosphonate (7.1 g, 33.2 mmol) was added to a solution containing 14.9 g (99.47 mmol) dry sodium iodide in 150 mL of dry acetone under nitrogen at 25 °C. Solutions were stirred overnight, after which the reaction mixture was evaporated to dryness. The solid mixture was triturated several times with ethyl acetate and the solvent was evaporated to dryness. The crude product was recrystallised using absolute ethanol to yield 5.1 g (70%), ^1^H NMR (500 MHz, D2O-DSS): δ = 8.10 (d, J 7.5 Hz, 2H), 7.61 (t, J 7.5 Hz, 1H), 7.50 (t, J 7.5 Hz, 2H), 3.55 (d, J 5.5 Hz, 3H). NMR spectrum of methyl benzoylphosphonate (MBP) was confirmed after the oxidation reaction, phosphonate signals were located in the 3.6 ppm region shifting 0.3 ppm from starting material as published by Brandt et al. (21).

For the inhibitor experiments, either oxamate or MBP were added to the medium at the beginning of the fermentation process to achieve a final concentration of 160 µg/L. In respiration experiments, inhibitors were added to a final concentration of 47 µg/mL in mid-exponential phase (OD620 ≈ 0.5).

### GC-MS analysis of intracellular and medium compounds

The pellets resulting from centrifugation were resuspended in 5 mL of Tris-HCl buffer at a pH of 8.2, and cells were lysed by sonication in a Sonics Vibra-Cell (Sonics and Materials Inc., Newton, CT). Organic compounds were extracted from the intracellular content using ethyl acetate and concentrated to dryness. Before GC-MS analysis, extracts were subjected to methylation in diethyl ether using N-nitroso-N-methylurea (NMU) in an alkaline medium. Extracellular volatile compounds were separated by liquid-liquid extraction using dichloromethane and then concentrated at 40 °C under nitrogen flow. Treatment of the samples and GC-MS analysis were performed as previously described by Martin et al. (28) using a Shimadzu-QP 2020 ULTRA mass spectrometer (Shimadzu Corp., Japan) equipped with a Stabilwax capillary column (30 m x 0.25 mm i.d., 0.25 µm film thickness; Restek, State College, PA).

### Identification and quantification

The analysed aromatic compounds were identified by comparing their linear retention indices using pure standards for 4-hydroxybenzaldehyde, 4-hydroxymandelic acid and 4-hydroxyphenylacetic acid. Mass spectra fragmentation patterns were also compared with those stored on commercial (29–31) and our own databases. Isotopically labelled compounds were quantified by ion fragment comparison. Benzoic acid was used as the internal standard for intracellular samples while 1-octanol was used for extracellular media.

### Coenzyme Q6 extraction and analysis by LC-MS

One milliliter of fermentation or respiration samples were centrifuged at 4 °C for 5 minutes, pellets were washed with distilled water. Then lipidic fraction was extracted as previously described by Ozeir et al., (2010). Briefly, 15-25 mg of pellet were resuspended in 50µl of KCl buffer (0,5mM), 25mM of coenzyme Q4 (Merk, Germany) was used as internal standard. Cells were disrupted using 600µL of methanol and 100mg of glass beads by vigorous vortexing for 10 minutes. Lipids were extracted with 800µL of pentanol in two steps, centrifuging 3 minutes at 1000g to recover the supernatant that was dried by N2 flux and resuspended in 100 µL of methanol.

Samples were analyzed by LC-MS. Chromatographic separation was performed on a reversed-phase column (Poroshell 120 EC-C18, 50 x 3 mm, 2.7 µ, Agilent), employing 5 mM ammonium acetate in H2O as mobile phase A and 5 mM ammonium acetate in methanol as mobile phase B. The elution program was as follows: 95% B for 2 min, followed by a rapid transition to 100% B, maintained for 4 min. The system was then re-equilibrated for 5 minutes. The flow rate was set at 800 µL/min, and the column temperature was maintained at 30 °C.

Mass spectrometric analysis was conducted on a QTRAP4500 (ABSciex) by monitoring Q6 in both its oxidized and reduced states, bearing either six 12C or six 13C atoms. Detection was achieved via ammonium adduct formation, with two specific transitions tracked per analyte. Fragmentation yielding the tropylium ion was utilized for quantification purposes [CoQ4ox m/z 472.2 /197.1 and 472.2 / 237.1; CoQ4red m/z 474.2 /197.1 and 474.2 / 237.1; CoQ6ox m/z 608.6 / 197.1 and 608.6 / 237.1; CoQ6red m/z 610.6 / 197.1 and 610.6 / 237.1; 13C₆-CoQ6ox m/z 614.6 / 203.1and 614.6 / 243.1; 13C₆-CoQ6red m/z 616.6 / 203.1and 616.6 / 243.1].

Quantification was performed by normalizing the peak area of each analyte to that of the corresponding Q4 internal standard. The sum of oxidized and reduced species was further normalized to the mass of the yeast pellet.

### Statistical analysis

ANOVA and post-hoc comparisons with Tukey’s test, using a significance value of 95%, were performed on 4-hydroxybenzaldehyde and Q6 concentrations resulting from fermentation and growth with S. cerevisiae strains. ANOVA comparisons were performed using STATISTICA 7.0 software (StatSoft, Tulsa, OK). Differences in mean concentrations of these compounds were evaluated using the least significant differences test.

## Acknowledgements

We thank Clarke Lab for suppling labelled ^13^C_6_-4-hydroxybenzoic acid and ^13^C_6_-*p*ABA . We wish to thank the following agencies for the financial support yielded to this work: CSIC Group Project #656 and CSIC Productive Sector Project #602 of UdelaR, Uruguay, Agencia Nacional de Investigación e Innovación (ANII) *Hanseniaspora vineae* FMV 6956 project. This work was also supported by the Programa de Alimentos y Salud Humana (PAyS) IDB - R.O.U. (4950/OC-UR) and a grant from Universidad de la República (Espacio Interdisciplinario, Centros 2020) to RR, and by the National Science Foundation Grant MCB-2343997 to CFC. Additional support obtained from Programa de Desarrollo de Ciencias Básicas (PEDECIBA, Uruguay).

We would also like to thank Lage y Cia-Lallemand Inc. Uruguay, and Oenobrands, France, for supporting our cooperation project in *H. vineae* metabolic characterization ANII ART_X_2022_1_172795.

---

**Table S1.** Nitrogen sources and amino acid composition used in the different media.

| Amino acid | *Growth*<br>(mg/L) | *Fermentation*<br>(mg/L) |
| --- | --- | --- |
| PRO | 60.3 | 120.6 |
| ALA | 12.1 | 24.2 |
| ARG | 90.4 | 180.8 |
| ASN | 18.1 | 36.2 |
| ASP | 42.2 | 84.4 |
| GLN | 24.1 | 48.2 |
| GLU | 60.3 | 120.6 |
| GLY | 6.0 | 12.0 |
| HIS | 143.5 | 161.2 |
| ILE | 24.1 | 48.2 |
| LEU | 536.2 | 572.4 |
| LYS | 530.1 | 560.2 |
| MET | 18.1 | 36.2 |
| PHE | 18.1 | 36.2 |
| SER | 48.2 | 96.4 |
| THR | 42.2 | 84.4 |
| TRP | 12.1 | 24.2 |
| TYR | 4.8 | 4.8 |
| VAL | 24.1 | 48.2 |
| Inorganic amonium source | (mg/L) | (mg/L) |
| DAP | 60.3 | 0 |

